# Potential of on-scalp MEG: Robust detection of human visual gamma-band responses

**DOI:** 10.1101/602342

**Authors:** Joonas Iivanainen, Rasmus Zetter, Lauri Parkkonen

**Author notes:** These authors contributed equally to this work (Joonas Iivanainen, Rasmus Zetter). Email addresses:* (Joonas Iivanainen), (Rasmus Zetter).

## Abstract

Electrophysiological signals recorded intracranially show rich frequency content spanning from near-DC to hundreds of hertz. Noninvasive electromagnetic signals measured with electroencephalography (EEG) or magnetoencephalography (MEG) typically contain less signal power in high frequencies than invasive recordings. Particularly, noninvasive detection of gamma-band activity (> 30 Hz) is challenging since coherently active source areas are small at such frequencies and the available imaging methods have limited spatial resolution. Compared to EEG and conventional SQUID-based MEG, on-scalp MEG should provide substantially improved spatial resolution, making it an attractive method for detecting gamma-band activity.

Using an on-scalp array comprised of eight optically-pumped magnetometers (OPMs) and a conventional whole-head SQUID array, we measured responses to a dynamic visual stimulus known to elicit strong gamma-band responses. OPMs had substantially higher signal power than SQUIDs, and had a slightly larger relative gamma-power increase over the baseline. With only eight OPMs, we could obtain gamma-activity source estimates comparable to those of SQUIDs at the group level.

Our results show the feasibility of OPMs to measure gamma-band activity. To further facilitate the noninvasive detection of gamma-band activity, the on-scalp OPM arrays should be optimized with respect to sensor noise, the number of sensors and inter-sensor spacing.

## 1. Introduction

Neuronal gamma-band (>30 Hz) synchronization seems a fundamental part of neural communication in the brain, having been linked to a multitude of cognitive functions, such as attentional selection (Tallon-Baudry et al., 1996; Fries et al., 2001) and working memory (Pesaran et al., 2002; Howard et al., 2003). The interest in electrophysiological gamma-band signals (the ‘gamma buzz’;Buzsaki 2006; Lachaux et al. 2012) was initiated by studies in the cat visual cortex that suggested that the binding of visual stimulus features might be mediated by neuronal synchronization at frequencies around 40 Hz (Eckhorn et al., 1988; Gray et al., 1989); the idea was then formulated as the binding-by-synchrony hypothesis (Singer, 1999). More recently, it has been also hypothesized that the precise timing of the gamma oscillations play a key role in information transfer between different cortical regions (Fries, 2015). While these hypotheses have assumed an oscillatory mechanism (usually present as a signal increase in a narrow frequency band; narrowband gamma), it is now appreciated that stimulation also usually induces a broadband increase in signal power in a range 40–150 Hz (broadband gamma or high-frequency activity; Crone et al. 2011; Lachaux et al. 2012) that is correlated with multi-unit activity, and thus reflects local cortical processing (Ray et al., 2008a; Manning et al., 2009; Ray & Maunsell, 2011).

Evidence of the link between gamma-band activity and cognitive functions was originally discovered in invasive experiments in non-human animals (e.g. Eckhorn et al., 1988; Gray et al., 1989; Fries et al., 2001). Similar findings were then reported in invasive human measurements using intracranial electroencephalography (iEEG) (e.g. Crone, 1998; Crone et al., 2001). Gamma-band activity has also been observed noninvasively in humans using scalp-EEG (e.g. Pfurtscheller & Neuper, 1992; Tallon-Baudry et al., 1996; Ball et al., 2008) and magnetoencephalography (MEG) (e.g. Adjamian et al., 2004; Hoogenboom et al., 2006). However, it is commonly understood that the noninvasive detection of gamma-band activity is difficult due to poor signal-to-noise ratio (SNR) and spatial resolution of the measurement methods (e.g. Dalal et al., 2009; Jerbi et al., 2009). Thus, the studies have usually relied on special stimuli crafted to maximize the gamma SNR.

The disparity in the SNR between invasively and noninvasively measured neural activity is dependent on the frequency: lower-frequency activity such as alpha (~10 Hz) and beta (~20 Hz) rhythms are typically equally present in noninvasive and invasive measurements, while higher-frequency activity is proportionally much weaker in noninvasive measurements (Jerbi et al., 2009). A likely reason for the low SNR of gamma-band signals detected by EEG/MEG is the short coherence length of the gamma sources, i.e. a patch of cortex producing synchronous gamma activity is typically much smaller than a patch producing synchronous activity at lower frequencies. Therefore, the key to detect gamma-band activity noninvasively is to improve the spatial resolution and sensitivity of the imaging method.

The spatial resolution of EEG is inherently limited by the spatial low-pass filtering of the neural electric field by the conductivity structure of the head (Pfurtscheller & Cooper, 1975; Srinivasan et al., 1998); the filtering leads to loss of spatial detail so that nearby sources cannot be readily separated easily or at all. By contrast, the neuromagnetic field measured by MEG is not as sensitive to head tissue conductivities (Hämäläinen et al., 1993; Stenroos & Nummenmaa, 2016; Iivanainen et al., 2017) so that the spatial detail of the field is mostly determined by the measurement distance. In conventional SQUID-based MEG, the distance of the sensors to head (at least about 2 cm) thus limits the spatial resolution.

Recent advances in the development of optically-pumped magnetometers (OPMs) (Budker & Romalis, 2007; Budker & Kimball, 2013) has enabled their use in MEG (e.g. Borna et al., 2017; Sheng et al., 2017; Boto et al., 2018; Iivanainen et al., 2019). OPMs, in contrast to SQUIDs, can be placed in very close proximity to the scalp, considerably boosting both spatial resolution and sensitivity to neural sources (Boto et al., 2016; Iivanainen et al., 2017). Thus, OPM-based *on-scalp* MEG shows a great promise both generally in the context of MEG and specifically for the detection of gamma-band activity.

In this work, we demonstrate that visual gamma-band responses can reliably be detected with on-scalp MEG based on currently-available OPMs at a similar or better SNR than with conventional SQUID-based MEG. Additionally, we demonstrate that source-level analysis of gamma activity can be performed with a limited number of channels only covering parts of the scalp.

## 2. Materials and methods

### 2.1. Subjects

Ten healthy volunteers (6 males, 4 females, 23–33 years of age, average 27.7 years) with no known history of neurological or psychiatric disorders participated in the study. The experimental design took into consideration the code of ethics as defined in the World Medical Association’s Declaration of Helsinki, and the study was approved by the Aalto University Ethics Committee. Informed consent was obtained from all participants.

### 2.2. Structural MRI acquisition and segmentation

T1-weighted structural MR images from previous studies were available for all subjects. The FreeSurfer software package (Dale et al., 1999; Fischl et al., 1999; Fischl, 2012) was used for pre-processing the MRIs and for segmentation of the cortical surfaces. For each subject, the surfaces of the skull and scalp were segmented using the watershed approach (Ségonne et al., 2004) implemented in FreeSurfer and MNE software (Gramfort et al., 2014). These surfaces were thereafter decimated to obtain three boundary element meshes (2562 vertices per mesh). For source estimation, the neural activity was modeled as a primary current distribution constrained to the surface separating the cortical gray and white matter and discretized into a set current dipoles (4098 locations per hemisphere, three orthogonal dipoles per location).

### 2.3. Experimental paradigm and stimuli

To evoke visual gamma-band activity, we used a stimulation paradigm originally presented by Hoogenboom et al. (2006) and thereafter employed in a multitude of studies (e.g. Hoogenboom et al., 2010; Scheeringa et al., 2011; Tan et al., 2016; van Pelt et al., 2018). In short, the experiment consisted of contracting sine-wave gratings projected on a screen in front of the subject inside a magnetically shielded room. In 90% of the trials, the contraction velocity of the grating increased at an unpredictable time but not earlier than 50 ms after the stimulus onset; the subject’s task was to detect this increase and report it as quickly as possible by the lifting the right index finger. The remaining 10% of the trials were a catch condition during which no velocity increase occurred. After each response, visual feedback of the correctness of the subject’s response was given.

The parameters of the stimulation paradigm were set same as in the work by Hoogenboom et al. (2006). The stimuli were projected on a semi-transparent back-projection screen using a projector (ET-LAD7700/L, Panasonic, Osaka, Japan; lens ET-D75LE3; refresh rate 60 Hz; resolution 1280× 1024), located outside the MSR. The distance from the eyes to the screen and the stimulus size were adjusted such that the outer diameter of the grating subtended a visual angle of 5°. Stimuli were presented using the ‘Presentation’ software package (Neurobehavioral Systems, Inc., Berkeley, CA, USA). Subject responses (finger lifts) were recorded using an in-house-built optically-triggered button.

During an approximately 1.5-hour session (including subject preparation), the experiment was performed using both OPM- and SQUID-MEG with each subject. The order of OPM and SQUID measurements was counterbalanced across subjects. Each subject completed one block of 100 trials per measurement, which lasted approximately 12 minutes. A one-minute resting-state measurement was performed after the primary task was completed.

### 2.4. MEG acquisition

OPM-MEG was recorded using an array of eight OPMs (Gen-1.0 QZFM, QuSpin Inc., Louisville, CO, USA). The OPMs were placed in a 3D-printed helmet with an identical geometry to that of a commercial 306-channel SQUID-MEG system (MEGIN Oy (formerly Elekta Oy), Helsinki, Finland). Individual OPMs were placed into sockets in the helmet, whose positions and orientations corresponded to those of the occipital sensors of the MEGIN system, and inserted until touching the head of the subject. The insertion depth was manually measured for each sensor. The helmet was attached to the chair the subject was seated in, and the subject’s head position inside the helmet was adjusted so that the OPM array covered the occipital cortex. To fix the position of the subject’s head inside the helmet, dummy sensors were inserted into sockets on the sides of the helmet so that they gently pressed on the head on each side. MEG–MRI co-registration was performed using an optical scanner as described by Zetter et al. (2019);an example of co-registered OPM positions with respect to the subject’s head is shown in Fig. 1. For Subjects 1 and 2, an additional ninth OPM was used (seen in Fig. 1, right side), however, for these subjects one OPM malfunctioned and was thus rejected from analysis. The OPM data (sensor bandwidth ~130 Hz) were recorded at 1-kHz sampling rate with an acquisition passband of 0.03–330 Hz using a data acquisition system based on the electronics of the commercial MEGIN system (Iivanainen et al., 2019). No additional magnetic shielding was used for the OPMs; the ambient-field amplitude and its drift inside our three-layer magnetically shielded room (MSR; Imedco AG, Hägendorf, Switzerland) was below ~10 nT and ~10 pT/hour, respectively, and thus did not pose any challenges for the operation of the OPMs (Iivanainen et al., 2019).

**Figure 1:**
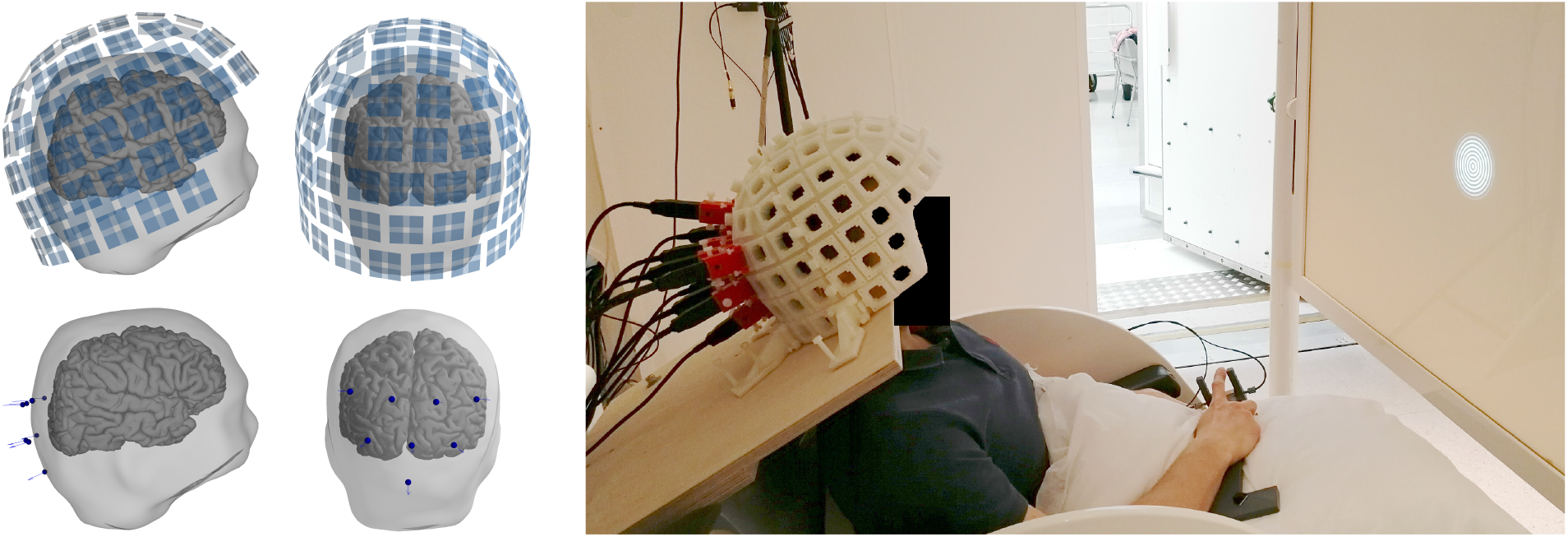
**Left:** SQUID (top) and OPM (bottom) sensor positions with respect to the head of a representative subject. **Right:** The OPM measurement setup in the magnetically shielded room.

SQUID-MEG was recorded using a whole-head 306-channel MEG system (Vectorview™ by MEGIN/Elekta Oy; 102 magnetometers; 204 planar gradiometers). The MEG signal was acquired with the same acquisition parameters as in OPM-MEG. MEG–MRI co-registration was performed using an electromagnetic digitizer in conjunction with head position indicator (HPI) coils. Approximately ~150 head-shape points were digitized and five HPI coils were applied in each subject. In both measurement modalities, 1 minute of data in the absence of a subject were also recorded.

### 2.5. Data analysis

We performed all MEG analysis using the MNE-Python software (master branch, checked out on 2018-10-19; Gramfort et al. 2014). Both OPM and SQUID data were bandpass-filtered to 0.1–130 Hz before further processing. Epochs were manually inspected and those containing visible artifacts were rejected.

#### Sensor-level analysis

Time–frequency representations (TFRs) of the responses were computed using Morlet wavelets. The frequency spacing for the TFR computation was 1 Hz, with the number of cycles for each frequency *f* set to *f*/2. The 500-ms period preceding stimulation was used as a baseline. The response data were divided by the mean of the baseline data, log-transformed and normalized by the standard deviation of the log-transformed baseline data.

Power spectral density (PSD) was computed using a multi-taper approach. Multiple orthogonal tapers (Slepian, 1978) were applied to each epoch, after which the spectral density of each channel was computed by averaging over each taper and epoch. The bandwidth of the multi-taper window function was set to 1 Hz. For baseline spectra, the 2.0–0.5-s period preceding stimulation was used. For the stimulation-period spectra, the time period 0.5–2.0 s following stimulation onset was used; the contribution of the initial evoked responses to the spectra was thus avoided. For quantitative analysis of the difference in spectral power between stimulation and baseline, we computed the ratio between the stimulation and baseline spectra for each sensor. Thereafter, we fitted a Gaussian function to the gamma-band peak in the relative spectrum of each sensor using the Trust Region Reflective algorithm as implemented in the *scipy* software library (Jones et al., 2001–). We estimated the gamma power increase (stimulation vs. baseline) by the peak amplitude of the Gaussian, the gamma frequency by the Gaussian peak frequency and the gamma bandwidth by the full width at half maximum (FWHM) of the Gaussian. The Gaussian function amplitude was constrained to be higher than 1, peak frequency was bounded to 40–70 Hz and the bandwidth to 1–20 Hz. The fitting was initialized with a guess amplitude of 10, center frequency of 52 Hz and a bandwidth of 8 Hz. In addition, the goodness-of-fits (GOFs) of the Gaussians were extracted. Sensors for which the fitted Gaussian function had GOF less than 0.5 or amplitude less than 1.05 were considered as not having a gamma response. This relative spectral power analysis was performed for all subjects for both OPM and SQUID data.

#### Source-level analysis

Forward models were computed using the MRI-derived three-shell boundary element models (see Section 2.2). OPMs were modelled as by Zetter et al. (2018), with eight integration points equally spaced within a 3-mm cube corresponding to the sensitive volume of the sensor. SQUIDs were modelled as in the MNE software. For the SQUID measurement, the 102 magnetometers and the entire whole-brain source space were used in the analysis. Due to the small number of OPM sensors, we restricted the source space in OPM analysis to cortical locations within 7 cm of the sensors, limiting the number of points in the source space to 2361 ±197 (mean±SD) across subjects. For the SQUID measurements, signal-space projection (SSP;Uusitalo & Ilmoniemi, 1997) based on an empty-room recording was applied to suppress artifacts in frontal channels that would otherwise corrupt the source estimates.

For source-level estimation of induced power, we employed DICS beamforming (Gross et al., 2001) as implemented in MNE-Python. To speed up the computation, data were decimated by a factor 3 before analysis. Cross-spectral density (CSD) matrices for baseline (2.0–0.5 s preceding stimulus onset) and stimulation (0.5–2.0 s following stimulus onset) time periods were computed within two frequency bands corresponding to alpha (7–13 Hz) and gamma bands (40–70 Hz). CSD computation was performed using a multi-taper approach similar to that used for PSD estimation, with the bandwidth of the multi-taper window function set to 1 Hz. The frequency bands were chosen on the basis of sensor-level TFRs as to encompass alpha- and gamma-band activity for all subjects. Based on the CSD matrices, separate spatial filters were computed for, and applied to, each time period and frequency band to create time- and frequency-specific source-power estimates. Diagonal loading regularization of 0.01 was applied when computing the spatial filters. The source orientation which produced maximal power was chosen, and no weight normalization was applied. Finally, the baseline-normalized difference in source power ((Stimulation–Baseline)/Baseline) was computed for both frequency bands, and across-subject grand averages were computed after morphing the data to the *fsaverage*’ template brain (Fischl et al., 1999). The grand-average source-level analysis was limited to those subjects for which clear gamma-band activity was present in the sensor-level analysis. A comprehensive tutorial of this type of analysis can be found in van Vliet et al. (2018).

## 3. Results

### 3.1. Sensor-level analysis

Examples of single-trial responses of all sensor types for Subject 6 are presented in Fig. 2. Gamma-band (40–70 Hz) power increases are visible in these responses. Across the sensor types, the responses are qualitatively similar. The OPM signal amplitude is larger than that of SQUID magnetometer by roughly a factor of two.

**Figure 2:**
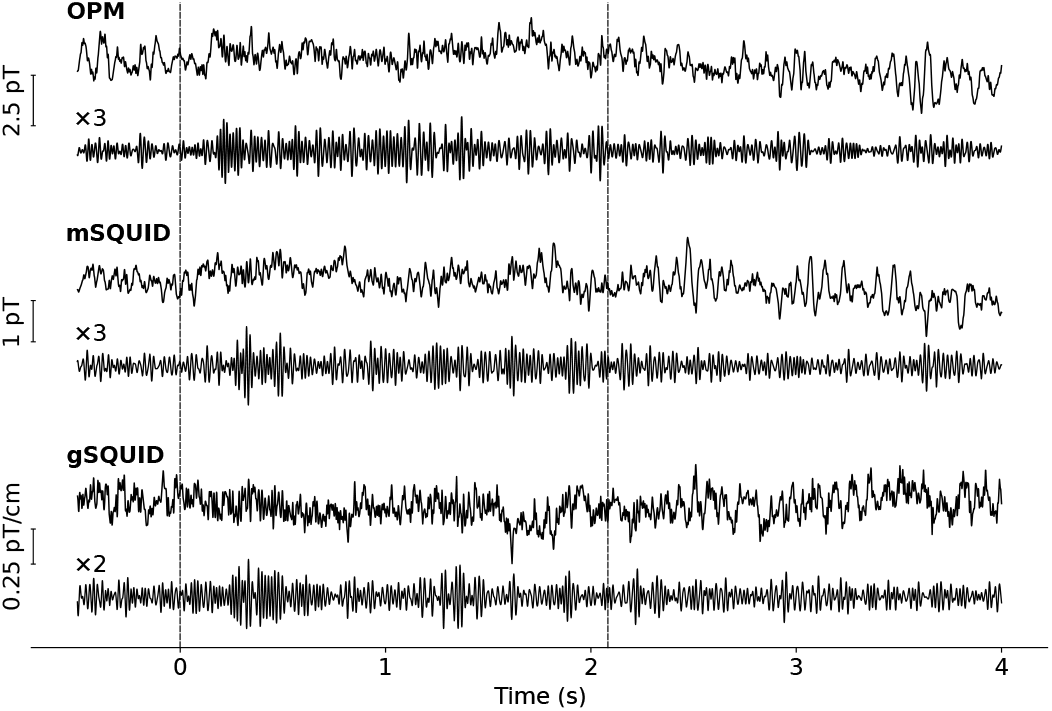
A representative single-trial response filtered to 0.1–130 Hz (upper traces) and to 40–70 Hz (lower traces) of OPM, SQUID magnetometer (mSQUID) and planar SQUID gradiometer (gSQUID) sensor with the largest gamma-band response (Subject 6). The dashed vertical lines indicate the onset and offset of the stimulus. Responses are from a trial without a change in the grating contraction velocity.

Averaged time–frequency representations (TFR) of the induced responses of OPM and SQUID magnetometers for Subject 6 are shown in Fig. 3A. The TFR of the sensor with the maximum gamma-band response amplitude is presented in Fig. 3B for each subject. Four out of the ten subjects (Subject 4, 5, 9 and 10) did not show clear gamma-band activity for either OPM or SQUID measurements while they did show a decrease in alpha–beta power with both sensor types. Another four subjects (Subject 2, 3, 6 and 8) demonstrated a simultaneous narrowband (30–40 Hz) and broadband (> 50 Hz) gamma increase.

**Figure 3:**
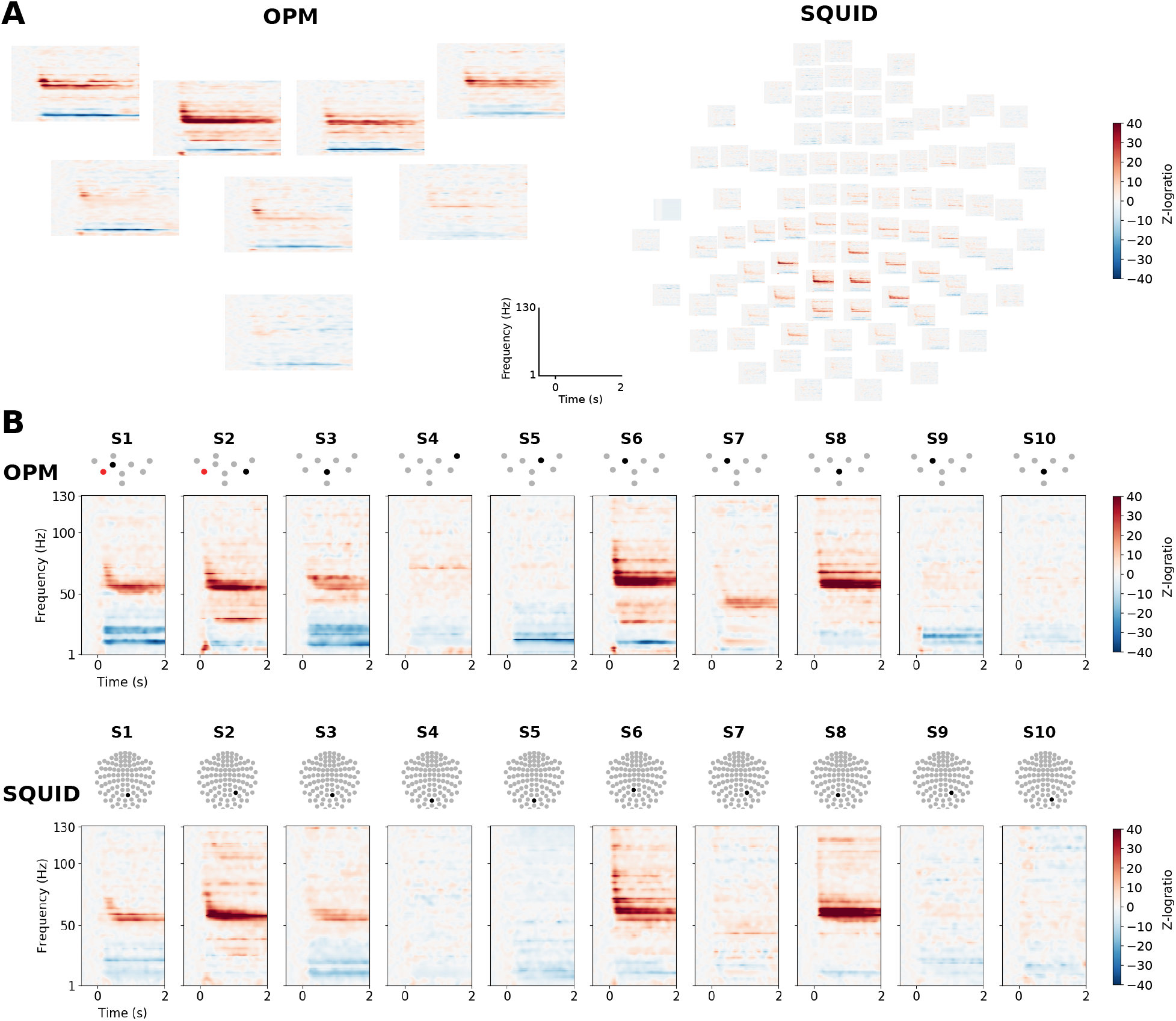
Time–frequency representations (TFRs) of the induced responses. **A:** Responses in a representative subject (Subject 6) in OPM (left) and SQUID (right) magnetometers. **B:** TFRs of the sensors with the maximal induced gamma-band response for OPMs (top row) and SQUID magnetometers (bottom row) across all subjects (one column per subject). The topographic sensor layouts indicate the sensor of the TFR (black) and a malfunctioning sensor (red) not included in the analysis.

The power spectral densities (PSD) and the metrics derived from them are presented in Fig. 4 for all subjects. As in the TFR analysis, the same four subjects did not show a considerable power increase within the gamma band for any sensor type. In Subject 7, the gamma power increase was present only in OPMs, while Subject 1 showed a gamma increase in both OPMs and SQUID magnetometers, but not in SQUID gradiometers. The goodness of the Gaussian fits to the PSDs of the sensors included in the analysis were 0.79±0.11, 0.84±0.07 and 0.82±0.08 for OPMs, SQUID magnetometers and SQUID gradiometers, respectively (mean±SD). For subjects with a significant gamma-band response at any sensor, the number of sensors showing this response was 3–8, 7–35 and 2–9 for OPMs, SQUID magnetometers and gradiometers, respectively. Relative to the baseline, the average gamma power across the subjects was 3.7, 2.5 and 2.0, while the maximum gamma power was 14.2, 13.1 and 4.1 for OPMs, SQUID magnetometers and gradiometers, respectively.

**Figure 4:**
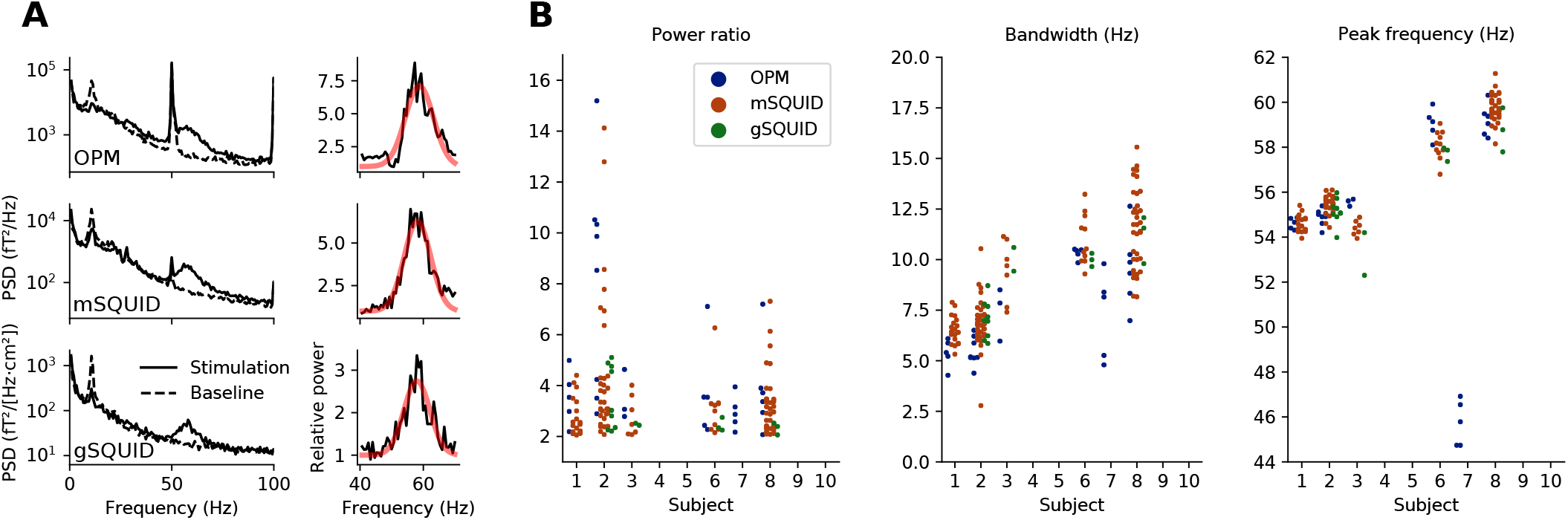
Power spectral analysis. **A: Left:** Power spectral densities (PSDs) estimated prior and during stimulation for the sensor with the highest gamma-band power for OPMs, SQUID magnetometers (mSQUID) and gradiometers (gSQUID) in a representative subject (Subject 6). **Right:** The ratio of the spectra (black) during stimulation and baseline, illustrating the power increase during stimulation, and the fitted Gaussian (red). **B:** Relative power (stimulation vs. baseline), bandwidth and peak frequency of the gamma response (extracted from the fitted Gaussian) across all subjects. The dots represent the values for individual sensors. Boxes correspond to the upper and lower data quartiles, and the horizontal line represents the median. Whiskers extend 1.5 times the interquartile range, and only data points past this range are shown individually.

### 3.2. Source-level analysis

Fig. 5 shows the average source-power difference between stimulation and baseline for both OPM and SQUID magnetometers across those six subjects (1, 2, 3, 6, 7, 8) for which clearly discernible gamma-band activity was present at the sensor level. Source estimates of individual subjects for both sensor types can be found in the Supplementary data (Figs. S1 and S2). The laterality of the group-level gamma-power estimate is the same in OPM and SQUID measurements while the laterality of alpha-power estimate differs between OPMs and SQUIDs. The shape and location of the high-amplitude region of the gamma-power (in yellow in Fig. 5) is slightly different between OPM and SQUID measurements.

**Figure 5:**
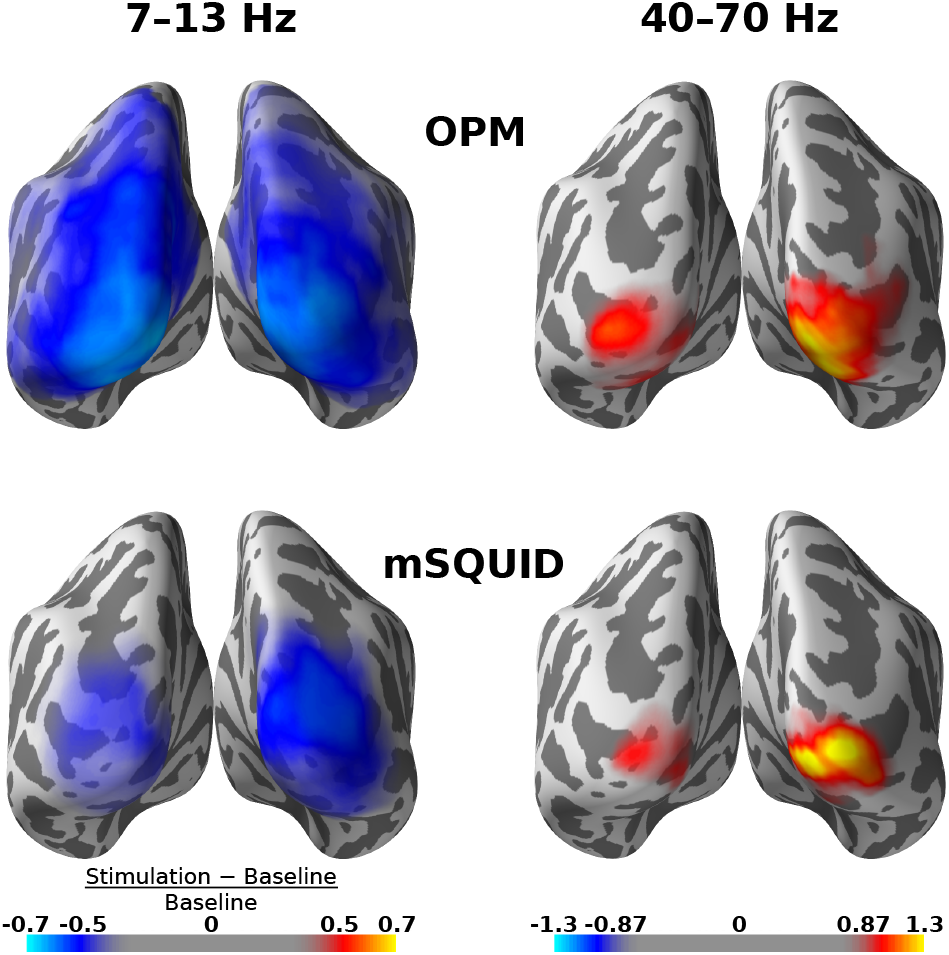
Grand-average normalized source-power difference between stimulation and baseline within alpha (7–13 Hz) and gamma (40–70 Hz) bands for SQUID magnetometers (top row) and OPMs (bottom row). The difference is visualized on the FreeSurfer ‘fsaverage’ template brain. Color maps are the same for both sensor types.

## 4. Discussion

In this work, we applied an experimental paradigm known to elicit gamma-band activity in the early visual cortices while measuring both OPM-based on-scalp MEG and conventional SQUID-based MEG. We showed that this type of gamma activity is well visible in on-scalp OPM-MEG as well as in conventional MEG, with gamma SNR (as compared to the baseline power) typically being higher in on-scalp OPM-MEG. In particular, we showed that with an 8-channel OPM array group-level source estimates of alpha and gamma power are comparable to those obtained with a 102-channel SQUID magnetometer array.

### 4.1. Gamma-band activity as measured by OPMs and SQUIDs

As Figures 3 and 4 demonstrate, gamma-band activity was seen with OPMs at least as well as with SQUID magnetometers. OPM signals have considerably higher absolute power than SQUIDs, as manifested already in the single-trial traces shown in Fig. 2 and in the PSDs of Fig. 4 due to the shorter distance between the OPMs and the neural sources.

Two subjects (Subject 2 and 6) displayed two separate peaks in the gamma frequency range (see Fig. 3); such a feature has also been observed earlier (Hoogenboom et al., 2006). In general, the gamma peak frequency varied considerably across subjects (Fig. 3 and 4), which has also been reported in several earlier studies (Hoogenboom et al., 2006, 2010; van Pelt et al., 2018) and which seems to show some genetic dependence (van Pelt et al., 2012) along with other factors such as gender and age (van Pelt et al., 2018). For some subjects, significant gamma-band activation was not detected in either OPM or SQUID measurements, at least not at the sensor level. Such a lack of visible gamma activity is also consistent with earlier literature, and may be due to many factors, such as differences in cortical folding, level of synchronous activity in cortical circuits, or even cognitive factors such as maintenance of the attentional focus in the stimulus.

### 4.2. Optimizing OPMs for detecting gamma-band activity

Due to the small number of sensors, the coverage of the cortex was limited in the OPM measurements. In addition, occipital areas were covered somewhat differently from subject to subject due to differences in the seating position and head shape; for some subjects, the OPM array was placed slightly too low to cover the whole occipital cortex adequately (see Supplements, Fig. S3). When using partial-coverage arrays and similar positioning methods as described here, specific care must be taken when adjusting the relative positions of the sensors to the head in order to achieve good coverage of the region of interest.

Our OPM array consisted of eight sensors with an intersensor spacing of approximately 3.4 cm, which is far from the ideal; the spacing should be roughly the distance from the sensors to the brain (Ahonen et al., 1993), which is about 1.5 cm measured from the scalp. Thus, to maximize the amount of spatial detail present at the sensors and thus the spatial resolution of the method, an on-scalp array with sensor spacing less than 1.5 cm covering the region of interest in the brain would be optimal. In the case of the occipital cortex, this would correspond to ~50 sensors.

The aforementioned limitations in sensor count and coverage and thus also in spatial sampling of the field limit our ability to fully utilize the benefits of on-scalp MEG arrays for modeling the neural sources. Nevertheless, we could successfully estimate the underlying sources from our OPM measurements, with the group-level estimates corresponding well to those based on the whole-head SQUID-magnetometer data (Fig. 5). The subject-specific estimates also appear fairly similar for those subjects that had visible gamma-band activity at the sensor level (see Supplementary data).

We have previously shown that the optical co-registration method we applied in the OPM measurement is sufficiently accurate for on-scalp MEG (Zetter et al., 2018, 2019). Thus, co-registration error should not be the limiting factor when improving OPM source estimates beyond those obtained from SQUID data.

Due to the physics of the OPM sensors, their bandwidth is limited in comparison to SQUIDs, which have no intrinsic maximum measurable frequency as far as MEG measurements are concerned. The bandwidth of the OPMs employed in this work extends to ~130 Hz (falling off 6 dB/octave), which is adequate for the narrowband gamma (40–70 Hz) we observed here. In contrast, broadband gamma in the visual cortex has been shown to extend at least to 200 Hz (Hermes et al., 2014). Thus, to study the entire gamma spectrum, the OPM bandwidth should be higher. In addition – besides broadband gamma – there are other high-frequency MEG signals beyond the current OPM bandwidth, e.g. axonal ~600-Hz ς-bursts (Curio, 2000) detected in S1 cortex in response to electric nerve stimulation as well as 150–250-Hz bursts linked to epileptic discharges (RamachandranNair et al., 2008; Zijlmans et al., 2012). As bandwidth and sensitivity are inherent trade-offs in OPMs (Budker & Romalis, 2007), further OPM sensor development is needed to meet the standards set by SQUIDs.

The current commercial OPMs can provide good-quality data, at least in environments where the ambient interference level is low, such as in our three-layer magnetically shielded room. Still, sensor noise (9–16 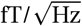 in our eight OPMs) rather than the 1/*f* background brain activity (Buzsaki, 2006; Miller et al., 2009) was limiting the SNR of the gamma-band responses. From the spectrum of background brain activity as measured with OPMs, we estimate that to make the OPM measurement limited by the brain background activity within the OPM bandwidth (~130 Hz), the sensor noise level should be below 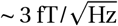.

Here, we used only eight OPMs to successfully record gamma-band activity. To study specific and localized brain activity with a good spatial resolution, such small arrays could be sufficient for many applications provided that the inter-sensor distance is short enough (< 1.5 cm). These arrays could be operated in a typical shielded room augmented with active shielding against the static (Boto et al., 2018) or static and dynamic (Iivanainen et al., 2019) components of the ambient field. The sensor arrays could also be operated in a low-cost person-sized magnetic shield (e.g. Borna et al., 2017). Thus, we expect that MEG could be adopted more widely with the availability of OPMs and compact shields as one does not need an expensive whole-scalp MEG system in spacious surroundings for every MEG application.

### 4.3. Narrow- vs. broadband gamma

There seem to be two or more separate phenomena that are commonly referred to as gamma-band activity: synchronized, narrowband gamma oscillations as well as broadband gamma spanning at least 50–150 Hz (Lachaux et al., 2012). Narrowband gamma, which has also been termed "binding gamma” within the visual system, can be observed at least in the visual (e.g. Hoogenboom et al., 2006), somatomotor (e.g. Bauer, 2006; Gross et al., 2007) and auditory systems (Brosch et al., 2002). There are several hypotheses concerning its role (e.g. Wang, 2010; Jensen & Mazaheri, 2010; Donner & Siegel, 2011; Fries, 2015). Broadband gamma, on the other hand, is associated with a general increase in locally synchronous neuronal firing regardless of stimulus modality and brain region (Mukamel, 2005; Liu & Newsome, 2006; Belitski et al., 2008; Ray et al., 2008b; Miller et al., 2014). Broadband gamma provides information about local processing in neuronal circuits and can thus serve as a highly focal marker of brain activity; for example, it can reveal spatial differences of the processing of highly similar stimuli (see e.g. Flinker et al., 2011). Whether and how narrow- and broadband gamma are related still remains an open question (Lachaux et al., 2012). Broadband gamma is often observed in invasive measurements (e.g. Crone et al., 2001; Edwards et al., 2005; Crone et al., 2006; Cervenka et al., 2013; Hermes et al., 2014; Bartoli et al., 2019) but rarely in noninvasive ones.

In this work, we studied mainly narrowband gamma oscillations (here 40–70 Hz) to compare OPM and SQUID recordings. The narrowband gamma elicited by the stimuli utilized in this work seems to stem from a large cortical area and be well visible in MEG. In contrast, the typical broadband gamma activity should have a short coherence length and thus be less visible in noninvasive measurements. We also analyzed the presence of higher-frequency activity (> 70 Hz) that should provide a proxy for the broadband gamma, see Fig. 3. Such high-frequency activity was discernible in some of the time–frequency representations but it did not stand out in the power spectra. At the source level, a slight power increase within the 70–130 Hz band is seen during stimulation (especially with SQUIDs); see Fig. S4. A more extensive analysis and a larger dataset would be needed for a more definitive and statistically rigorous comparison of broadband activity as measured by OPMs and SQUIDs.

As on-scalp MEG should have better spatial resolution than conventional SQUID-based MEG (Boto et al., 2016; Iivanainen et al., 2017), the use of on-scalp MEG to measure broadband gamma is attractive and could open a new noninvasive window into the functioning of the human brain. The noninvasive detection and localization of broadband gamma would be greatly facilitated by more sensitive OPM sensors assembled into arrays that provide dense spatial sampling of the neuromagnetic field and a good coverage of the cortical area of interest.

## Conclusions

We have shown here that visual gamma-band responses can be measured with a small on-scalp OPM array with response quality comparable to that obtained with a conventional whole-scalp SQUID-based MEG. Gamma power and signal-to-noise ratio were larger in OPMs compared to SQUIDs. To further facilitate the noninvasive detection of gamma-band activity, the on-scalp OPM arrays should be optimized with respect to sensor noise, the number of sensors and inter-sensor spacing.

## Acknowledgments

Research reported here was supported by the European Research Council under ERC Grant Agreement no. 678578, the National Institute of Neurological Disorders and Stroke of the National Institutes of Health under Award Number R01NS094604 the Finnish Cultural Foundation under grant nos. 00170330 and 00180388 (JI) and the Instrumentarium Science Foundation under grant no. 180043 (RZ). The content is solely the responsibility of the authors and does not necessarily represent the official views of the funding organizations.

