## Supplementary data for "Potential of on-scalp MEG: Robust detection of human visual gamma-band responses"

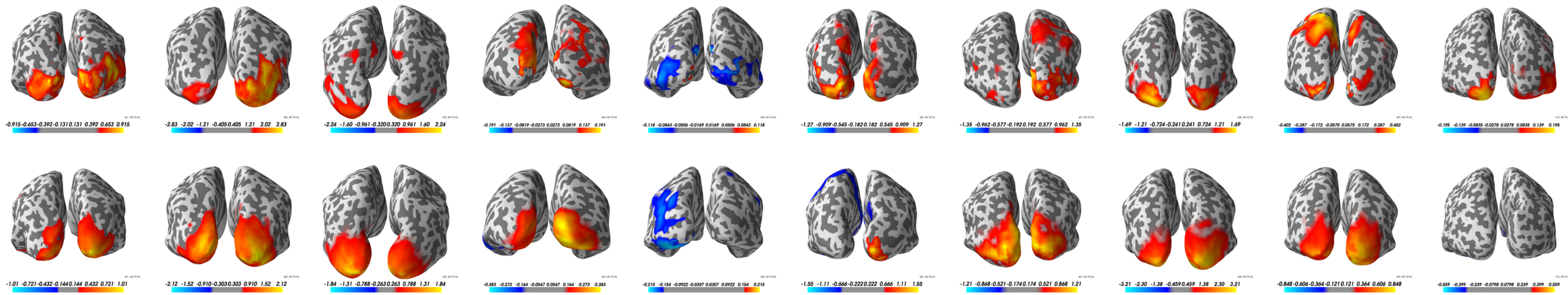

Figure S1: Subject-specific gamma-band source estimates for OPMs (top row) and SQUID magnetometers (bottom row). Color scales are specific to each estimate.

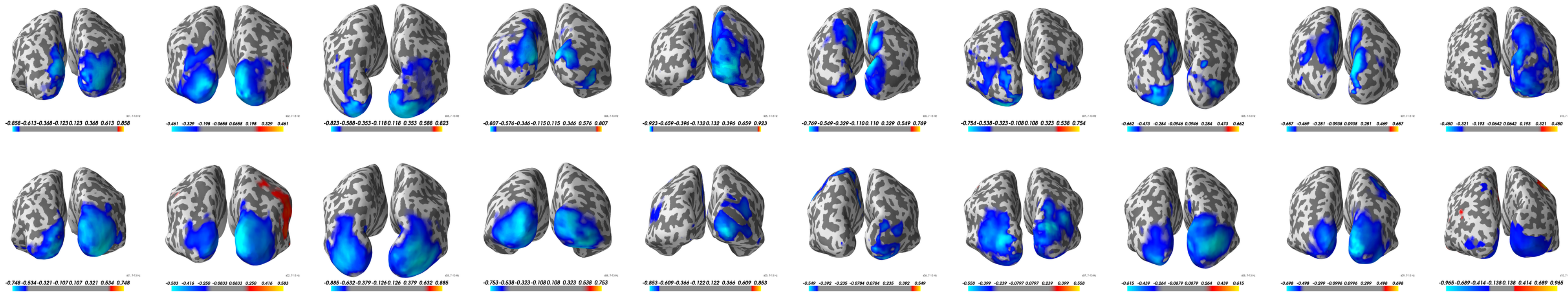

Figure S2: Subject-specific alpha-band source estimates for OPMs (top row) and SQUID magnetometers (bottom row). Color scales are specific to each estimate.

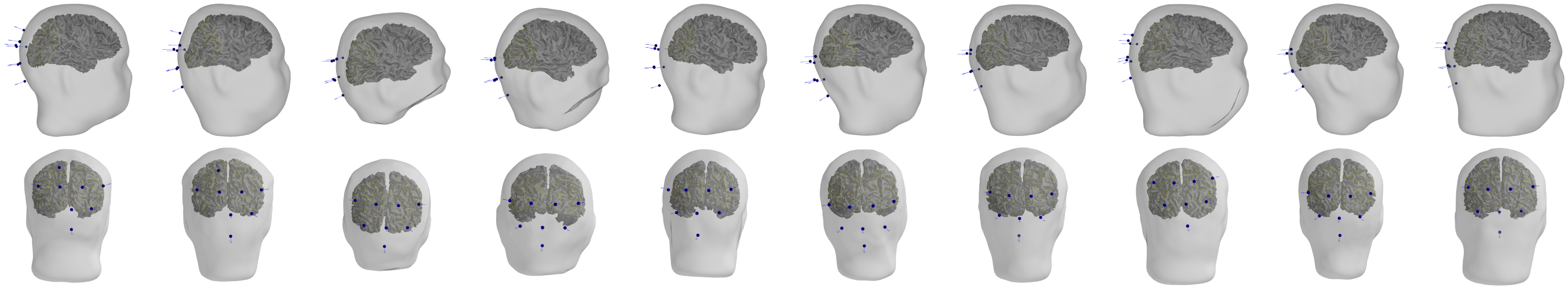

Figure S3: OPM sensor positions in all subjects; lateral (top row) and occipital views (bottom row). The source space (yellow points) was constrained on the surface separating the white and gray matter.

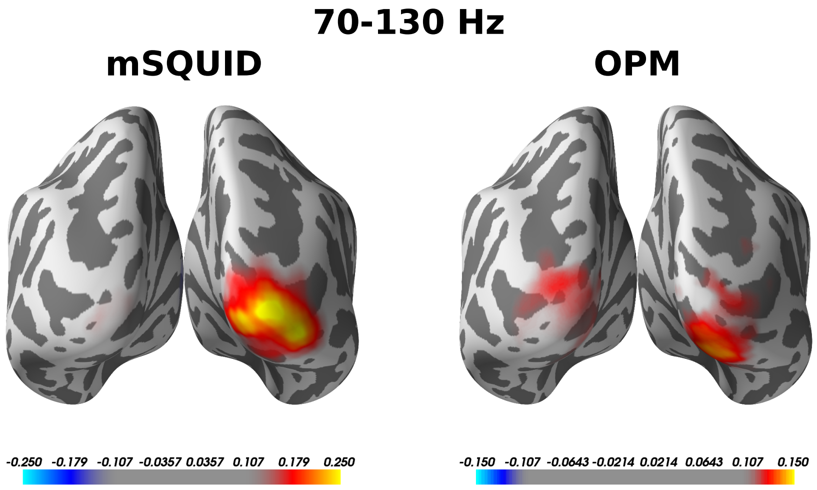


Figure S4: Grand-average baseline-normalized source power difference between stimulation and baseline within 70–130-Hz band for SQUID magnetometers (left) and OPMs (right).
